# Effect of visual cues in addition to moderate auditory cues on temporal coordination: A comparative study in humans and chimpanzees

**DOI:** 10.1101/290379

**Authors:** Lira Yu, Masaki Tomonaga

## Abstract

Our previous studies reported that chimpanzees share an ability to produce spontaneous temporal coordination with humans (Yu & Tomonaga, 2015; 2016). However, it remains unclear how visual cues of an interacting partner’s movement influence on the emergence of tempo convergence. The current study conducted a comparative study in humans and chimpanzees under the same experimental setup as that used in Yu & Tomonaga (2016). Three conditions, including baseline, paired-invisible and paired-visible, were prepared. In the baseline condition, the participants produced the repetitive tapping movement alone. In contrast, in the other two paired conditions, the participants in a pair produced the tapping movement concurrently while facing a conspecific partner. However, in the paired-invisible condition, a visual barrier was placed in between the participants to control visual cues of an interacting partner’s movement. Moderate auditory cues, corresponding to each participant’s tapping movement, were presented throughout the conditions. Results showed that there are significant changes on the tapping tempo between baseline and the paired-invisible condition, whereas there are little changes between paired-invisible and paired-visible condition in both species. The current stepwise analysis across three conditions demonstrates that auditory cues were more influential than additive visual cues of an interacting partner’s movement on the tempo convergence in humans and chimpanzees.

## Introduction

Our two previous studies revealed that chimpanzees have an ability to produce spontaneous temporal coordination when they interact with another conspecific individual. In the first study conducted under the side-by-side setup, we demonstrated that one chimpanzee gradually matches her tapping movement with auditory cues which was from an interacting partner (Yu & Tomonaga, 2015). In the second study conducted under the face-to-face setup, we found a general tendency on temporal coordination in chimpanzees (Yu & Tomonaga, 2016). These findings indicate that auditory cues can solely facilitate an interaction between chimpanzees but additional visual cues do enhance the influence of an interacting partner’s movement on production of one’s own movement. However, it remains unclear how additional visual cues play a role in spontaneous temporal coordination in chimpanzees.

In the current study, we examined an effect of visual cues of an interacting partner’s movement on one’s own spontaneous tapping movement in chimpanzees. Moreover, we tested human participants under the same experimental setup to directly compare with the findings from the chimpanzees.

## Methods

### Participants

We tested two pairs of chimpanzees (*Pan troglodytes*), in total four chimpanzee individuals (ID Number of Great Ape Information Network is from https://shigen.nig.ac.jp/gain/, see Watanuki et al., 2014). One pair was from kin-relationship, 34-year-old mother [Chloe](GAIN-ID C-0441) and her 14-year-old daughter [Cleo] (GAIN-ID C-0609). Both chimpanzees joined our previous experiments (Yu & Tomonaga, 2015; 2016). Another chimpanzee pair was from non-kin relationship, 37-year-old female Pendensa [Pen] (GAIN-ID C-0095) and 38-year-old female [Mari] (GAIN-ID C-0274). All the four chimpanzees lived together with seven other chimpanzees in an enriched outdoor compound with attached indoor residences in Primate Research Institute, Kyoto University (Matsuzawa, 2006). All the chimpanzees have had extensive experience with perceptual and cognitive studies (Tomonaga, 2001; Matsuzawa, 2003; Matsuzawa et al., 2006). In humans, we tested two pairs from non-kin relationship. One human pair consists of 24-year-old female [H1X] and 60-year-old female [H1Y]. Another human pair consists of 24-year-old female [H2X] and 46-year-old female [H2Y]. The human participants were acquainted with each other because they were graduate students or staff members in the institute. The care and use of the chimpanzees were carried out in accordance with the 3rd edition of the Guide for the Care and Use of Laboratory Primates issued by Primate Research Institute, Kyoto University in 2010. The experimental protocol for the chimpanzees was approved by the Animal Welfare and Animal Care Committee of the same institute and the Animal Research Committee of Kyoto University (2012-043, 2013-040 and 2014-037). The research protocol for human participants was reviewed and approved by the Human Ethics Committee in Primate Research Institute, Kyoto University (2015-02).

### Apparatus

LED push buttons (Fuji Electric, 2 × 2 cm) were used to produce repetitive and rhythmic finger tapping movements from each participant. The two buttons were aligned horizontally 12.5 cm apart in a transparent acrylic panel. The distance between the two acrylic panels was approximately 24 cm. In other words, the distance between two participants in a pair was at least 24 cm. An automated universal feeder was used to deliver a food reward to the chimpanzees. All the equipment and experimental events were controlled by a computer located outside the experimental booth.

### Task

Each trial began with illumination of two push buttons. The participants were required to tap either one of the two buttons. When one button was pressed, its light turned-off and the participants were again required to tap the remaining lighted-up push button. When this button was also pressed, its light turned-off and another button in the other side illuminated with 0 ms delay. The participants were required to tap the illuminated button with her own spontaneous tapping tempo until a trial Finish. A trial finished after the alternative button pressing from 25 to 45 times. The chimpanzees received a piece of apple in the end of each trial. The inter-trial interval was 2 seconds.

### Conditions

Three conditions were prepared: baseline, paired-invisible and paired-visible condition. In the baseline (BL) condition, the participants produced the tapping movement alone. In contrast, in the other two conditions, two participants in a pair produced the tapping movement concurrently while facing each other. These two conditions were prepared for visibility control. In the paired-invisible (Pi) condition, black-colored paper was placed between two acrylic panels so that visual cues of an interacting partner’s movement can be blocked from each other (Figure 1a). On the other hand, in the paired-visible (Pv) condition, the black occluder was removed and visual cues of an interacting partner’s movement was available between participants (Figure 1b). Throughout the three conditions, there was a mechanical click sound (approximately 50 dB) from the button pressing.

**Figure 1.**
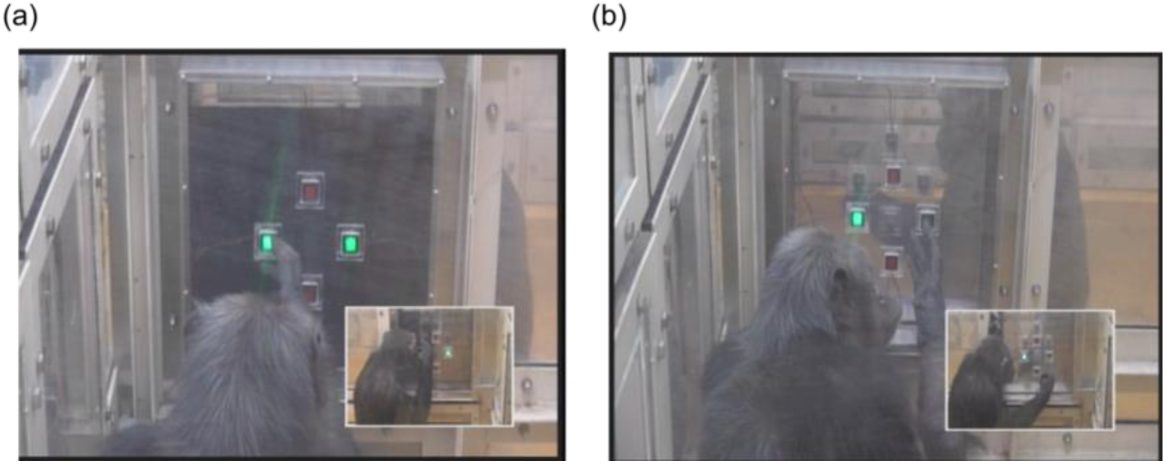
A pair of the chimpanzees (Chloe and Cleo) conducting the finger-tapping task under two test conditions. (a) Tapping in the Paired-invisible condition where the occluder prevents visual cues of the partner’s movement. (b) Tapping in the Paired-visible condition where the tapping movement is visible from each other.

### Procedure

Both chimpanzees and humans were tested under the same experimental setup and schedule. To measure one’s own spontaneous tapping tempo, the participants were first conducted the tapping task independently. This baseline (BL) condition was conducted for two consecutive days. Each day consisted of 5 blocks of six trials. Each trial required the participants to tap from 25 to 45 times.

After the two consecutive days for the baseline, the participants conducted the task concurrently with their conspecific partner. The two test conditions were conducted under ABBA schedule (A: Paired-visible, B: Paired-invisible). Only one condition was conducted in a day, and it was conducted for two consecutive days. Thus, the experimental days for each test condition were 4 days in total. Each day consisted of 5 blocks of six trials, and each trial required 25 to 45 taps. During these test conditions, the timing of a trial Start and Finish was occurred at the same time between two participants in a pair. In every each block, either one of the participants in a pair was randomly designated to finish a trial with producing alternative tapping from 25 to 45 times.

### Analysis

Tapping intervals, which is an inverse form of tempo, was calculated from the alternative tapping movements. Few very short (<100 ms) and long (>2 s) tapping intervals were removed as outliers for each participant. The tapping intervals were averaged for each block (n=10 for BL; n=20 for Pi and Pv, respectively).

Two sample *t*-tests assuming unequal variance were conducted in stepwise. Firstly, to confirm an effect of auditory cues from an interacting partner’s movement, the mean tapping intervals between baseline (BL) and paired-invisible (Pi) were compared for each participant and for each pair. Next, to examine an effect of additional visual cues of an interacting partner’s movement, the mean tapping intervals between paired-invisible (Pi) and paired-visible (Pv) were compared. No Bonferroni or Holm correction was conducted because the current comparisons were planned independently.

## Results

The mean of tapping intervals in the three conditions, including baseline (BL), the paired-invisible (Pi) and paired-visible (Pv), were examined for each participant in all four pairs in chimpanzees and humans (Figure 2). Firstly, the comparisons between BL and Pi revealed that all the participants, except for one chimpanzee (Cleo, *t*(24) = 1.88, *p* = 0.072), show a significant difference in their tapping intervals between the two conditions (Chloe, *t*(27) = 3.80, *p* < 0.001; Pen, *t*(17) = 11.05, *p* < 0.001; Mari, *t*(27) = 4.65, *p* < 0.001; H1X, *t*(27) = 18.93, *p* < 0.001; H1Y, *t*(14) = 13.10, *p* < 0.001; H2X, *t*(14) = 3.54, *p* = 0.003; H2Y, *t*(27) = 5.71, *p* < 0.001). Secondly, the comparisons between Pi and Pv revealed that five out of eight participants show no significant difference between the conditions (Cleo, *t*(31) = 1.04, *p* = 3.02; Pen, *t*(35) = 0.19, *p* = 0.848; H1X, *t*(30) = 0.78, *p* = 0.439; H1Y, *t*(26) = 0.42, *p* = 0.68; H2X, *t*(37) = 0.32, *p* = 0.747). The remaining three participants showed a similar pattern in that they produce faster tapping movement in the paired-visible condition in comparison with the paired-invisible condition (Chloe, *t*(35) = 2.15, *p* = 0.038; Mari, *t*(37) = 3.3, *p* = 0.002; H2Y, *t*(37) = 4.22, *p* < 0.001). These findings suggest that auditory cues were more influential than additive visual cues of an interacting partner’s movement on spontaneous tapping movement in both species.

**Figure 2.**
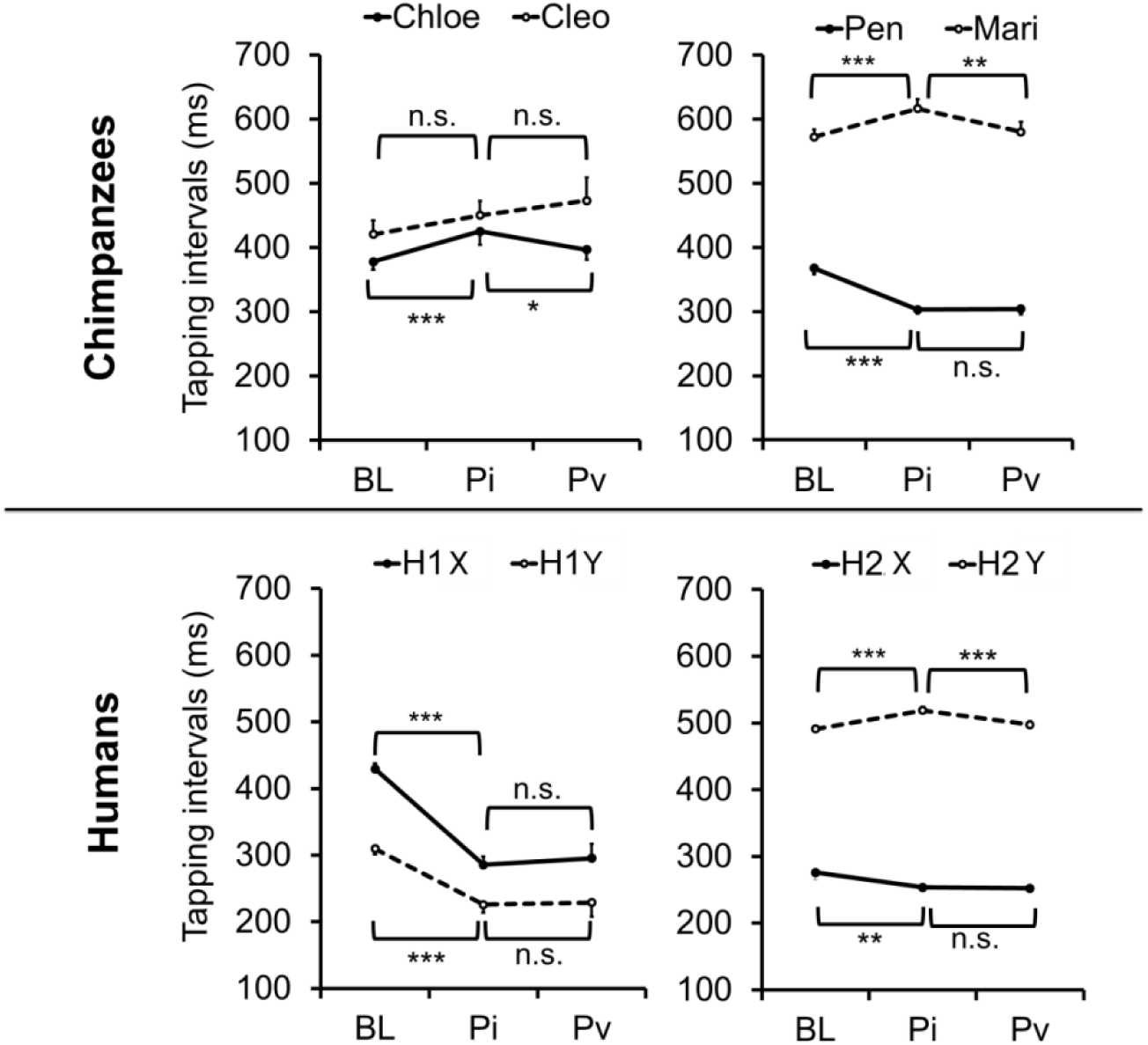
Mean of the tapping intervals under three conditions (baseline, Paired-invisible and Paired-visible) for each participant in each pair in chimpanzees and humans. Error bars indicate 95% confidence intervals of the mean.

To examine in what direction did the changes in tapping intervals occur, a ratio of the mean tapping intervals between the participants was calculated for each pair (relatively slow tapping individual’s tapping interval was divided by the fast tapping individual’s tapping intervals, see Figure 3). Both chimpanzees and humans showed the ratio which is close to 1:1 or 2:1 in Pi condition in comparison with BL. Moreover, the comparisons between Pi and Pv showed that the ratio in the Pv condition slightly deviated from that of the Pi condition. The results especially from the Pi condition demonstrate that there are two different types of the tempo convergence in both chimpanzees and humans.

**Figure 3.**
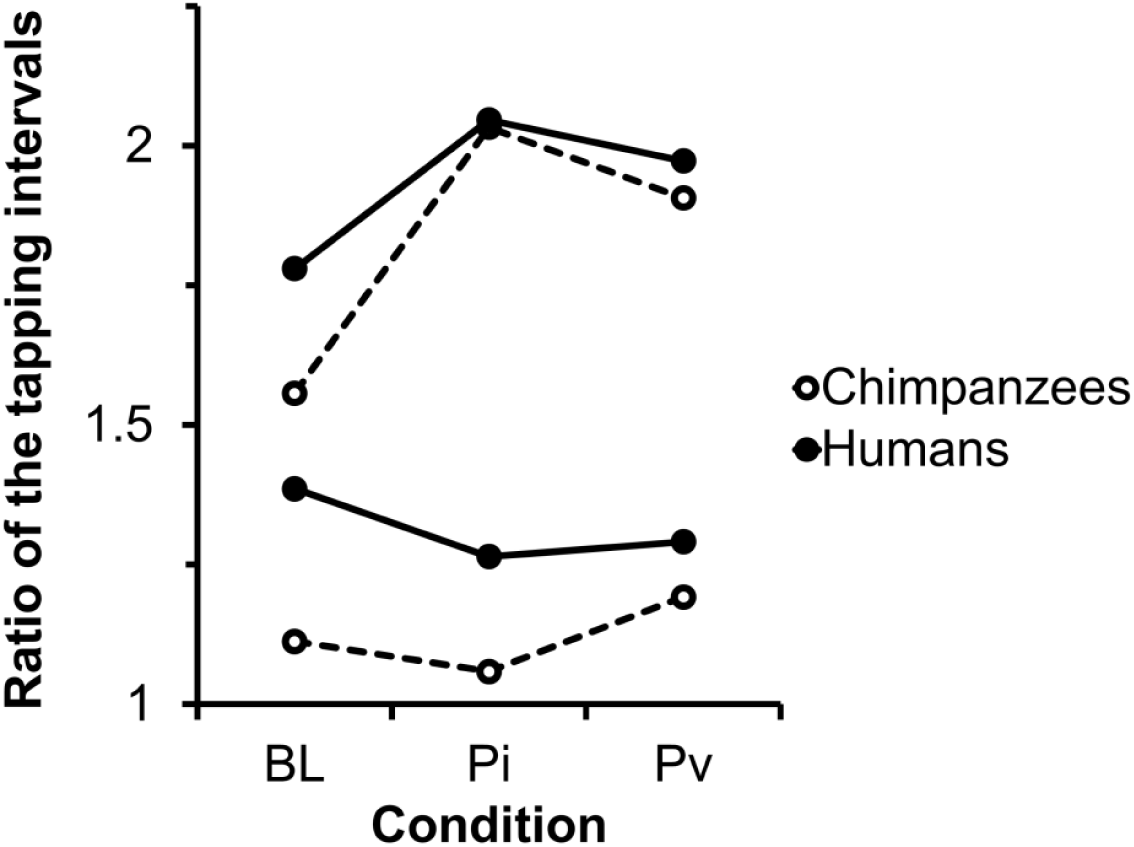
Each dot represents a ratio of the mean tapping intervals between two participants in each pair for each condition. The dots connected by lines represent data from the same pairs.

## Discussion

The current study investigated the causal effect of multimodal cues of an interacting conspecific partner’s movement on spontaneous tempo convergence in chimpanzees and humans. The results demonstrate that visual cues have little effect on the tempo convergence but auditory cues play a critical role in the tempo convergence in both species. This finding supports our previous study (Yu & Tomonaga, 2015), reporting that one chimpanzee spontaneously show the tapping alignment to her partner’s tapping movement even only auditory cues of the partner’s movement were available. Moreover, this finding is consistent with the previous studies in humans (Repp & Penel, 2002; 2004), reporting that humans are more strongly attracted to auditory rhythms than to visual rhythms.

The ratio of the tapping intervals between the participants in each pair indicate that the tempo convergence fall into either 1:1 or 2:1 match in both species. Previous studies in humans reported that polyrhythms (such as 3:2 or 5:3) are less stable than simple rhythms in which one frequency is an integer multiple of the other (such as 1:1, 2:1 or 3:1), and thus often resulting in transitions to simple rhythms (Fraisse, 1982; Treffner & Turvey, 1993). Therefore, the current results suggest that chimpanzees and humans may have a shared form of the stable tempo convergence.

In summary, the current study demonstrates that chimpanzees are similar with humans in terms of that auditory cues are sufficient to facilitate tempo convergence. Moreover, both species show harmonic tempo convergence when they concurrently produce the rhythmic tapping movement with an interacting partner.

## Acknowledgments

We thank T. Matsuzawa, M. Hayashi, I. Adachi and Y. Hattori, and the staff of the Language and Intelligence Section and Center for Human Evolution Modeling Research of Primate Research Institute, Kyoto University for their support and daily care of the chimpanzees. This study was financially supported by the Grants-in-Aid for JSPS fellows (#244525 and #16F16001), JSPS-MEXT KAKENHI (#15H05709, #16H06283, #20002001, #23220006, #24000001), JSPS-CCSN, Global COE programs (A06, D07) and the JSPS Leading Graduate Program in Primatology and Wildlife Science (U04) at Kyoto University.

